# Brain-derived neurotrophic factor derived from peripheral sensory neurons plays a critical role in pain chronification

**DOI:** 10.1101/147165

**Authors:** Shafaq Sikandar, Michael S. Minett, Joanne Lau, Queensta Millet, Sonia Santana-Varela, John N Wood, Jing Zhao

## Abstract

Multiple studies support the pro-nociceptive role of brain-derived neurotrophin factor (BDNF) in pain processes in the peripheral and central nervous system. We have previously shown that nociceptor-derived BDNF is implicated in inflammatory pain. Microglial-derived BDNF has also been shown to be involved in neuropathic pain. However, the distinct contribution of primary afferent-derived BNDF to chronic pain processing remains undetermined. In this study, we used Advillin-CreERT2 mice to delete *Bdnf* from all adult peripheral sensory neurons. Conditional BDNF knockouts were healthy with no sensory neuron loss. Behavioural assays and *in vivo* electrophysiology indicated that spinal excitability was normal. Following formalin inflammation or neuropathy with a modified Chung model, we observed normal development of acute pain behaviour, but a deficit in second phase formalin-induced nocifensive responses and a reversal of neuropathy-induced mechanical hypersensitivity during the later chronic pain phase in conditional BDNF knockout mice. In contrast, we observed normal development of acute and chronic neuropathic pain in the Seltzer model, indicating differences in the contribution of BDNF to distinct models of neuropathy. We further used a model of hyperalgesic priming to examine the contribution of primary afferent-derived BDNF in the transition from acute to chronic pain, and found that primed BDNF knockout mice do not develop prolonged mechanical hypersensitivity. Our data suggest that BDNF derived from sensory neurons plays a critical role in mediating the transition from acute to chronic pain.

## Introduction

Brain-derived neurotrophic factor (BDNF), a member of the neurotrophin family, acts as a modulator of neuronal excitability and synaptic plasticity (Pezet and McMahon, 2006). We have previously shown that nociceptor-derived BDNF regulates the excitability of spinal neurons and plays a crucial role in inflammatory pain (Zhao *et al.*, 2006), supporting other studies that have demonstrated neurotrophin regulation of synaptic transmission and long term plasticity in pain pathways (Kerr *et al.*, 1999; Slack *et al.*, 2004; Melemedjian *et al.*, 2013). Although multiple studies support the role of BDNF expressed throughout the nervous system in nociceptive processing in the peripheral and central nervous system (Merighi *et al.*, 2008; Ferrini and De Koninck, 2013; Nijs *et al.*, 2015a), the distinct contribution of primary afferent derived BDNF to chronic pain processing remains undetermined. Here we demonstrate the critical role of BDNF expressed in peripheral sensory neurons in the transition from acute to chronic pain.

BDNF expressed in dorsal root ganglia (DRG) is released in an activity-dependent fashion in the spinal dorsal horn to activate trkB receptors on second order neurons or primary afferent endings (Thompson *et al.*, 1999; Lever *et al.*, 2001; Chen *et al.*, 2014). It has long been known that exogenous BDNF can facilitate spinal reflexes and increase primary afferent evoked post synaptic currents (Kerr *et al.*, 1999; Garraway *et al.*, 2003). BDNF derived from Na_v_1.8-expressing nociceptive neurons contributes to inflammatory pain induced by intraplantar carrageenan and intramuscular NGF, but does not affect the development of neuropathic pain in a modified Chung model (Zhao *et al.*, 2006). In contrast, BDNF derived from microglia drives pain behaviour in a model of sciatic nerve cuffing (Coull *et al.*, 2005), and BDNF derived from myelinated afferents is dramatically increased following spinal nerve transection (Obata and Noguchi, 2006a; Obata *et al.*, 2006a). However, a pro-nociceptive role of BDNF derived from sensory neurons alone is yet to be determined. Some studies using models of hyperalgesic priming have demonstrated a role of BDNF in mediating persistent pain in migraine and following acute inflammation (Melemedjian *et al.*, 2013; Burgos-Vega *et al.*, 2016), which suggest that BDNF expression overall can drive chronic pain. However, whether BDNF released from sensory neurons alone can mediate the transition of acute to chronic pain is still unknown.

Developmental effects or compensatory mechanisms with embryonic gene deletion can mask the normal role of genes and expression of phenotypes in an adult system. This may be relevant to genes, such as *Bdnf*, which are expressed in different cellular populations and have temporal changes in expression throughout development (Cohen-Cory *et al.*, 2010; Kasemeier-Kulesa *et al.*, 2015). This can be addressed with the inducible CreERT2 system for better spatial and temporal deletion of genes (Feil *et al.*, 2009). In this study, we deleted *Bdnf* from all sensory neurons in adult animals by crossing an Advillin-CreERT2 strain (Lau *et al.*, 2011) with floxed *Bdnf* mice. We determined the contribution of sensory neuron-derived BDNF on acute pain processing and pain chronification using inflammatory, neuropathic and hyperalgesic priming models of chronic pain.

## Materials and Methods

### Transgenic mice

Homozygous floxed *Bdnf* mice (*Bdnf*^fl/fl^) carrying *lox*P sites flanking exon 5 in the *Bdnf* gene (Rios *et al.*, 2001) were interbred with Advillin-CreERT2 mice (Lau *et al.*, 2011) expressing a tamoxifen-inducible modified Cre recombinase under the control of the peripherally-restricted sensory neuron-specific advillin promoter to obtain *Bdnf*^fl/fl^; Avil-CreERT2 mice. Genotyping of mice for *Bdnf*^fl/fl^ and Advillin-CreERT2 was performed by standard PCR and the following primers were used (5’-3’): Bdnf1 (forward) GCCTTCATGCAACCGAAGTATG and Bdnf2 (reverse) TGTGATTGTGTTTCTGGTGAC; Advillin1 (forward) CCCTGTTCACTGTGAGTAGG and Advillin2 (reverse) AGTATCTGGTAGGTGCTTCCAG; and Advillin-CreERT2 (reverse) GCGATCCCTGAACATGTCCATC. The expected sizes of amplicons are: floxed *Bdnf* – 487 bp; *Bdnf* wildtype – 437 bp; Advillin-CreERT2 – 180 bp; Advillin wildtype – 480 bp. To delete the *Bdnf* gene, *Bdnf*^fl/fl^; Avil-CreERT2 mice were treated with tamoxifen at the age of 8 – 12 weeks as described (Lau *et al.*, 2011). At the same time, the littermate *Bdnf*^fl/fl^ control mice received the same treatment in parallel. Ten days after the tamoxifen treatment (5-day intraperitoneal injection, 2mg per day), *Bdnf*^fl/fl^; Avil-CreERT2 knockout mice and *Bdnf*^fl/fl^ control mice were used for experiments.

All animal assays performed in this study were approved by the United Kingdom Home Office Animals (Scientific Procedures) Act 1986. Both female and male mice aged 8–14 weeks were kept on a 12-h light/dark cycle and maintained under standard conditions (21 ± 1 °C, food and water *ad libitum*).

### Immunohistochemistry

Three mice from each group (*Bdnf*^fl/fl^; Avil-CreERT2 and *Bdnf*^fl/fl^) were used for DRG cell counting. After CO_2_ euthanasia, L4 DRGs were excised and rapidly frozen with O.C.T compound on dry ice and cut in serial sections (11 μm thick). Every 8th section through the entire DRG was collected onto superfrost slides and dried at room temperature for 4 hours. Immumohistochemistry and cell counting were performed as previously described (Zhao et al., 2006). Briefly, after three washes in phosphate-buffered saline (137 mm NaCl, 10 mM Na_2_HPO_4_, 1.8 mM KH_2_PO_4_, 2.7 mM KCl, pH 7.4) containing 0.3% Triton X-100 (PBST), sections were incubated with 10% goat serum in PBST at room temperature for 1 hour and then incubated with primary antibodies, mouse anti-peripherin (1:1000; Sigma, catalogue #P5117) and rabbit anti-N200 (1:200; Sigma, catalog #N4142), overnight at 4°C. After three washes in PBST, sections were incubated with secondary antibodies, goat anti-rabbit Alexa Fluor 488 antibody (1:1000, Invitrogen, catalog #A-11017) and anti-mouse IgG Alexa Fluor 594 (1:1000; Invitrogen, catalog #A-11037) for 2 hours in dark. After 3 washes in PBST, the sections were mounted with VECTASHIELD HardSet Antifade Mounting Medium (Vector, catalog #H-1400) and visualised using a Leica DMRB microscope, a Hamamatsu ORCA-R2 digital camera and HCIamge 2.0.1.16 software. The sample images were analysed using the cell counter plugin for ImageJ. Every visible cell was counted whether the nucleolus was present or not. Each DRG was counted twice, and the results were pooled. The percentages of NF200-positive, peripherin-positive and double-stained cells were calculated for each section. Mean and SEM of these percentages were evaluated for wild-type and mutant groups and significance was determined using a two-tailed unpaired heteroscedastic *t*-test.

### *In vivo* Electrophysiology

Electrophysiological recordings were performed by an experimenter blind to genotype (*Bdnf*^fl/fl^; Avil-CreERT2, *n* = 86 neurons) and their littermate controls (*Bdnf*^fl/fl^, *n* = 105 neurons). Mice were anaesthetized with isofluorane (4%; 0.5L/min N2O and 1.5L/min O2) and secured in a stereotaxic frame. Anaesthesia was reduced and maintained at 1.5% isoflurane for the remaining duration of the experiment. A laminectomy was performed to expose L3–L5 segments of the spinal cord and extracellular recordings were made from Wide-dynamic Range (WDR) neurons in the deep dorsal horn (lamina III–V, 200–600 μm) using parylene-coated tungsten electrodes (A-M Systems, USA) in *Bdnf*^fl/fl^ controls and *Bdnf*^fl/fl^; Avil-CreERT2 knockout mice. Mechanical and thermal stimuli were applied to the peripheral receptive field of spinal neurons on the hind paw glabrous skin and the evoked activity of neurons was visualised on an oscilloscope and discriminated on a spike amplitude and waveform basis using a CED 1401 interface coupled to Spike2 software (Cambridge Electronic Design, UK). Natural stimuli (dynamic brush, vF hairs 0.16g – 26g, noxious prod 100 and 150 g cm^−2^ mechanical stimulation; thermal water jet 35 – 45 °C) were applied in ascending order of intensity to receptive fields for 10 s and the total number of evoked spikes recorded. Statistical significance for differences between littermates and *Bdnf*^fl/fl^; Avil-CreERT2 mice was determined using a 2 way repeated measures ANOVA with Bonferroni post-tests for all measures.

### Mouse behaviour

All behavioural experiments were performed by an experimenter blind to genotype (*Bdnf*^fl/fl^ *n* = 8, *Bdnf*^fl/fl^; Avil-CreERT2 *n* = 10). Mechanical nociceptive thresholds were measured using Randall-Selitto apparatus (Ugo Basile) that applies pressure to the tail with a 3-mm^2^ blunt conical probe using a 500g cutoff. Mechanical sensitivity of the inferior half of the abdomen was assessed using von Frey hair application as described previously (Minett *et al.*, 2014). Rotarod testing was performed over 5 minutes with the initial starting ramp increasing from 4rpm to 40rpm over 30s using a mouse-adapted apparatus (IITC). Thermal nociceptive thresholds were determined by measuring hind paw withdrawal latency using the Hargreaves apparatus (IITC) at a ramp of 1.5°C s^−1^ with a 30s cutoff and latency for nociceptive behaviour on a hot plate (Ugo Basile) at 50°C and 55°C. Thermal place preference plates (Bioseb) were used to determine temperature preference of mice to 30°C (warm), 14°C (cool) and 5°C (cold) temperatures versus a room temperature 20°C plate. All place preference thermal tests were performed with reversed plate temperature settings and the average of the two responses used for statistical analyses. Nociceptive behaviour to cooling acetone was measured following application of one acetone drop to the hind paw and nociceptive behaviours measured for 1 minute. Statistical significance for differences between littermates and BDNF knockouts was determined using a *t*-test.

### Formalin test

The two groups of animals (*Bdnf*^fl/fl^ *n* = 6, *Bdnf*^fl/fl^; Avil-CreERT2 *n* = 6) were singly housed in Perspex boxes and allowed to habituate to the testing environment for 30 minutes. Animals were then injected with formalin (15μl i.pl., 5% dilution of stock formalin (40% w/v) in saline) in the hind paw. Nociceptive behavior was measured as licking and biting of the injected paw only. Nociceptive behaviour was recorded at 5 minute intervals for a duration of 60 minutes and divided in 2 phases, the first phase lasting 0 – 10 minutes and the second phase 10–60 min. Statistical significance for differences between littermates and *Bdnf*^fl/fl^; Avil-CreERT2 mice over the time course or for comparison of biphasic behaviour was determined using 2 way repeated measures ANOVA with Bonferroni post-tests and *t*-test, respectively.

### Neuropathic pain models

Neuropathic pain was assessed with two peripheral nerve injury models. For the partial sciatic nerve ligation model (Seltzer model), the surgical procedure was carried out as previously described (Zhao *et al*, 2010). Briefly, animals were anaesthetised using isoflurane. A 0.5 cm incision was made in the skin of the upper left leg and blunt scissors were used to dissect muscle layers apart to access the sciatic nerve. A tight ligation between ½*-*⅓ of the nerve was made using 6–0 mersilk suture (Ethicon, UK). The skin was closed using 4–0 mersilk sutures (Ethicon, UK). For spinal nerve ligation, a modified version of the Kim and Chung model (Kim and Chung, 1992) of peripheral neuropathy was adapted for use on mice (Minett *et al.*, 2012). Briefly, under the same anesthesia, a midline incision was made in the skin of the back at the L4-S2 levels and a further incision through the left paraspinal muscle was made to access the transverse processes at the L4-S1 levels. The L4 transverse process was removed using a blunt fine forceps and the left L5 and L4 spinal nerves identified. The L5 spinal nerve was separated and tightly ligated with 8–0 silk sutures and transected just distal to the ligature. The incision was closed in layers.

Statistical significance for differences between *Bdnf*^/fl^ littermate controls and *Bdnf*^/fl^; Avil-CreERT2 knockout mice was determined using a 2 way repeated measures ANOVA with Bonferroni post-tests. Statistical significance of changes in pain behaviour over time compared to baseline within each group was determined with a 1 way repeated measures ANOVA with Dunnett’s post-tests.

### Hyperalgesic priming model

A model of hyperalgesic priming was first established with wildtype male C57BL/6J mice. Mechanical sensitivity of the hind paw was measured using the up-down method to determine the 50% mechanical withdrawal threshold to von Frey application (Chaplan *et al.*, 1994). The primed group of mice was administered with carrageenan (i.pl. 1% 20μl, Sigma Aldrich) on Day 0, followed by administration of Prostaglandin E2 (PGE2) in the same paw (i.pl. 100ng in 25μl, Cayman Chemical) in both primed and unprimed mice on Day 6 (*n* = 6 in both groups). We tested *Bdnf*^fl/fl^; Avil-CreERT2 male mice (*n* = 6) and their littermate male controls *Bdnf*^fl/fl^ (*n* = 7) in this model of hyperalgesic priming using the same time course of drug administration. Statistical significance for differences between primed/unprimed mice or *Bdnf*^fl/fl^; Avil-CreERT2/littermates was determined using a 2 way repeated measures ANOVA with Bonferroni post-tests. Statistical significance of changes in pain behaviour over time compared to baseline within each group was determined using a 1 way repeated measures ANOVA with Dunnett’s post-tests.

### Statistical analysis

All values are presented as Means ± SEM. Data were analysed using the GraphPad Prism 7. Student’s *t*-test (two-tailed) was used for comparison of difference between two groups. Multiple groups were compared using one-way or two-way analysis of variance with Dunnett’s or Bonferroni post-tests, respectively. Differences were considered significant at *P* < 0.05.

## Results

### Generation of sensory neuron-derived BDNF knockout mice

To generate *Bdnf*^fl/fl^; Avil-CreERT2 mice, we crossed *Bdnf*^fl/fl^ mice with Avil-CreERT2 mice on a *Bdnf*^fl/fl^ background. The genotypes of offspring were analyzed with standard PCR. The result shows that homozygous a floxed *Bdnf* band and a Avil-CreERT2 band appeared in BDNFconditional knockouts (Fig. 1A). In contrast, the *Bdnf*^fl/fl^ littermate control mice only have a homozygous floxed *Bdnf* band (Fig. 1A). We have previously confirmed *Bdnf* deletion in DRGs of BDNF knockout mice using Real-Time qRT-PCR, showing about 70% reduction of BDNF mRNA in DRG 10 days after tamoxifen injection (Neumann *et al.*, 2016). The remaining ~30% of BDNF mRNA may come from satellite glial cells (Wetmore and Olson, 1995), and/or may be attributed to degrading/degraded mRNA in DRG neurons.

**Figure 1.**
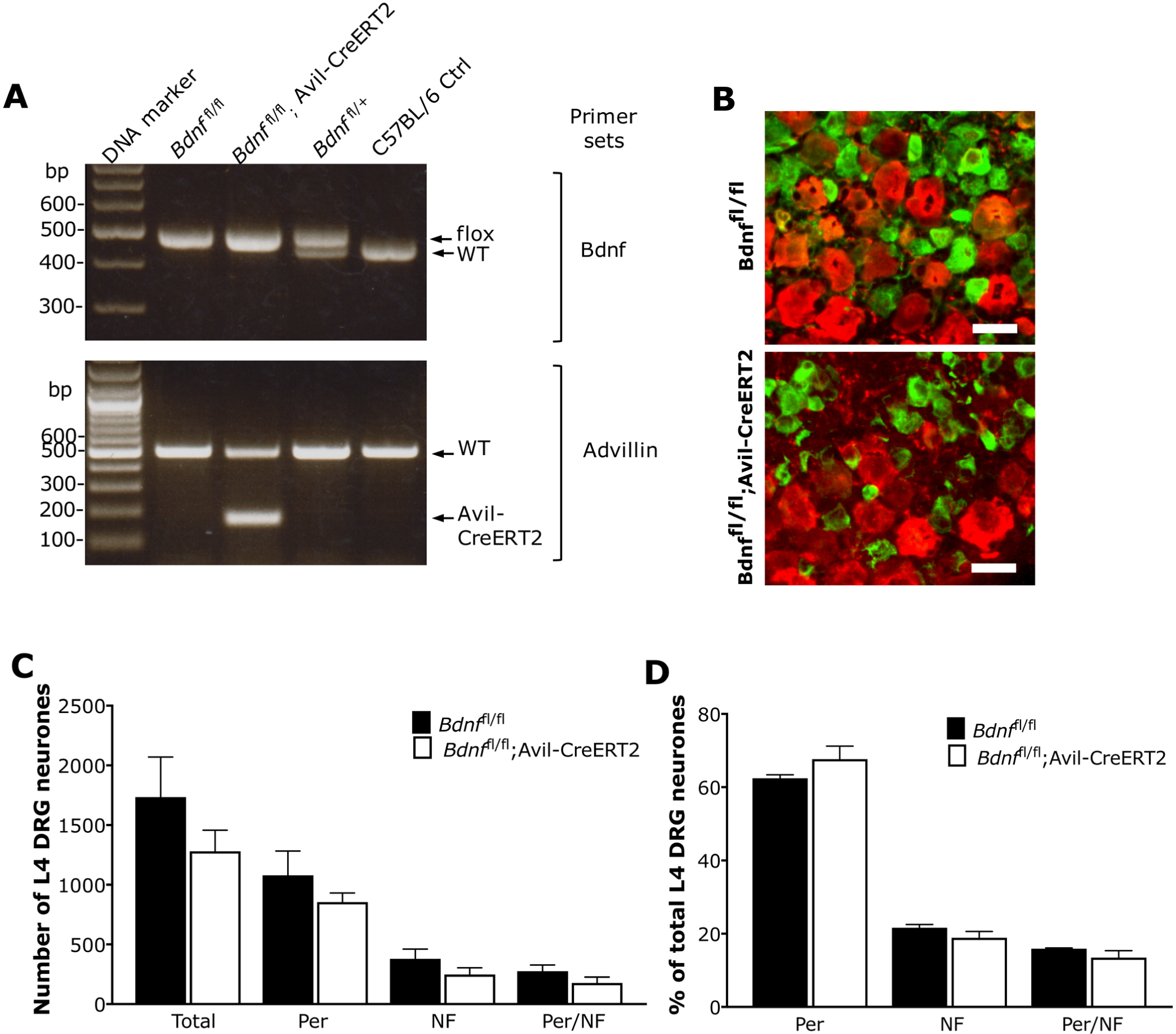
Characterization of BDNF knockout mice. (**A**) Genotyping analysis with PCR. The representative gel analysis of PCR products using both the BDNF primer set and the Advillin-CreERT2 primer set is shown in top and bottom panels, respectively. Mice homozygous for the floxed *Bdnf* band and heterozygous for the Advillin-CreERT2 band were defined as *Bdnf*^fl/fl^; Avil-CreERT2 mice. Mice only homozygous for the floxed Bdnf band were defined as *Bdnf*^fl/fl^ littermate controls. The genomic DNA either from C57BL/6J mice or from heterozygous floxed Bdnf (*Bdnf*^fl/+^) mice was used as controls. (**B**) DRG sections were labelled with large diameter DRG neuron marker neurofilament (in red), and small-medium diameter DRG neuron marker peripherin (in green). Scale bar, 50 μm. (**C**) The total number of DRG neurons expressing neurofilament (NF) and peripherin (Per) in L4 DRG sections were counted. (**D**) The proportions of neurofilament and peripherin expressing neurons calculated. Both total number and proportion are normal in BDNF knockout mice (*n* = 3) compared to littermate controls (*n* =3). Data were analysed with Student’s *t*-test and *P* > 0.05.

### Conditional BDNF deletion in adult mice does not affect the survival of DRG neurons

We then performed immunohistochemical staining of lumbar DRG sections to determine whether deletion of BDNF from sensory neurons affects survival or population of DRG neurons using small to medium diameter neuron (nociceptor) marker peripherin, and large diameter neuron marker neurofilament heavy chain (NF200). Our data show that most nociceptors were labelled with anti-peripherin, and most large diameter DRG neurons were NF200 positive in both *Bdnf*^fl/fl^; Avil-CreERT2 mice and *Bdnf*^fl/fl^ mice (Fig. 1B). There was no apparent difference between *Bdnf* conditional knockouts and littermate controls in the total number and proportion of neurofilament and peripherin-positive neurons (Fig. 1C and 1D). This is similar to our previous findings with BDNF deletion from Nav1.8-expressing neurons (Zhao *et al.*, 2006).

### Acute pain processing

To assess sensory coding of spinal neurons, we used extracellular recordings of evoked activity of L4 deep dorsal horn wide dynamic range neurons in *Bdnf*^fl/fl^ mice and *Bdnf*^fl/fl^; Avil-CreERT2 mice. Mechanical and thermal stimuli were applied to the peripheral hind paw receptive field of neurons and evoked action potentials were recorded as previously described (Sikandar *et al.*, 2013; Minett *et al.*, 2015). For mechanical sensory coding, we compared evoked activity of dorsal horn neurons to a range of intensities of punctate vF hairs (Fig. 2A), as well as two noxious prods and a low threshold brush stimulation applied to the whole receptive field of the hind paw (Fig. 2D). For thermal sensory coding, we compared evoked activity to innocuous and noxious heat (Fig. 2B), as well as noxious cold with ethyl chloride (Fig. 2C). Dorsal horn neurons in both conditional BDNF knockouts and littermate controls showed graded coding to increasing intensities of both mechanical and thermal stimuli. However, we observed no significant difference between groups of evoked activity to any mechanical or thermal stimulation parameter (*P* > 0.05 in all measures, 2 way repeated measures ANOVA with Bonferroni post-tests). These findings indicate that deletion of *Bdnf* from dorsal root ganglia does not alter normal spinal sensory coding of mechanical and thermal stimuli.

**Figure 2.**
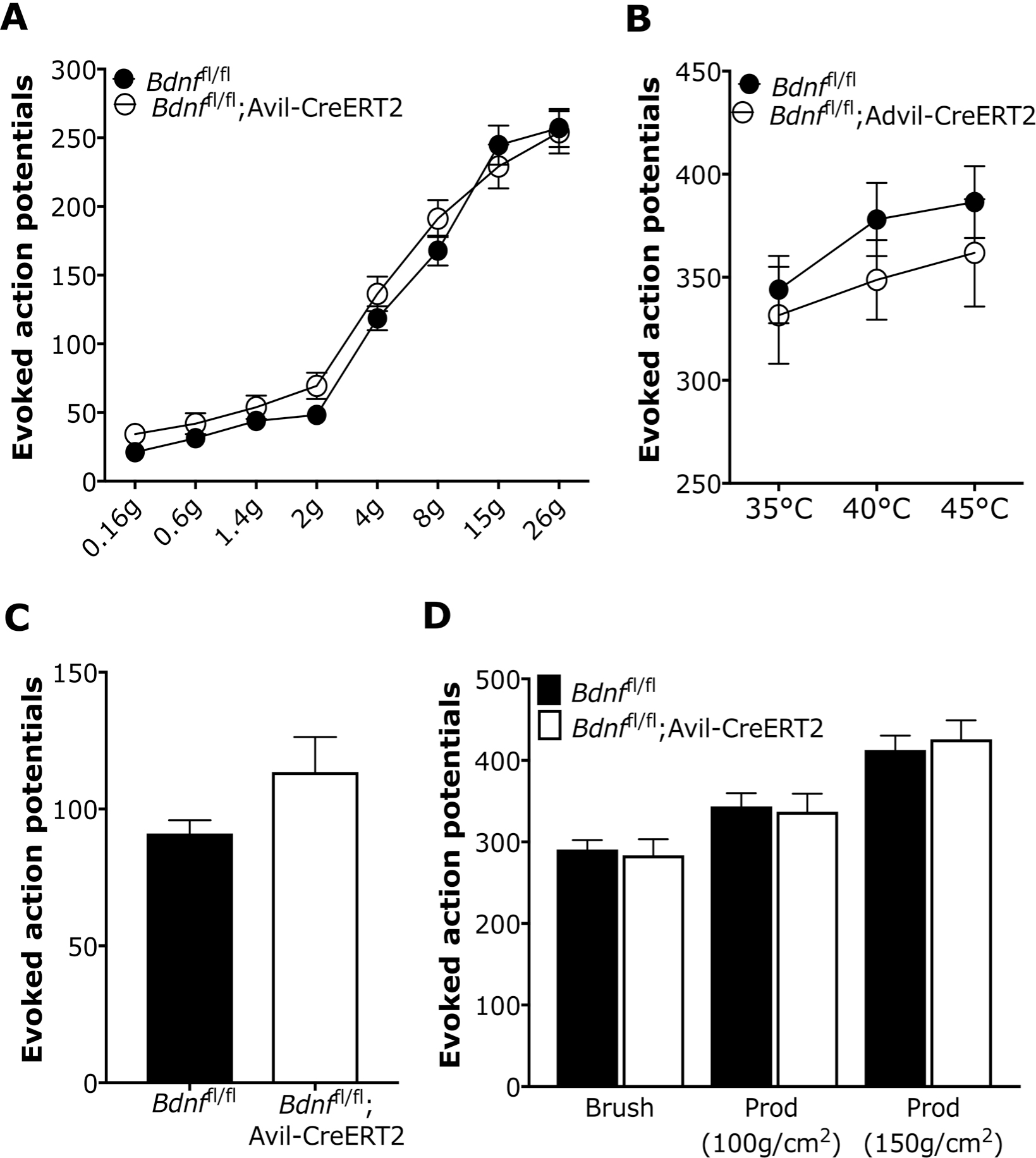
Evoked activity of Wide-dynamic Range neurons in deep dorsal horn was assessed by *in vivo* electrophysiology. (**A**) Evoked activity to mechanical punctate stimulated with von Frey hair on hind paw. (**B**) Thermal stimuli. (**C**) Noxious cold. (**D**) dynamic brush and prod stimulation. 86 WDR neurons from *Bdnf*^fl/fl^; Avil-CreERT2 and 105 WDR neurons from *Bdnf*^fl/fl^ micewere recorded. Data were analysed with 2 way repeated measures ANOVA with Bonferroni post-tests and *P* > 0.05 in all measures.

Standardised behavioural assays were used to asses thermal and mechanical pain thresholds in BDNF knockout mice (Fig. 3). Deletion of *Bdnf* from sensory neurons had no impact on motor function, as shown by normal rotarod activities measured in speed (Fig. 3A, *t*-test, *P* = 0.21), time spent on the rotarod (Fig. 3B, *t*-test, *P* = 0.26) and distance travelled (Fig. 3C, *t*-test, *P* = 0.18). We also observed no significant difference in mechanical pain thresholds using the Randall-Sellitto test with a noxious probe applied to the paw (Fig. 3D, *t*-test, *P* = 0.72) and tail (Fig. 3E, *t*-test, *P* = 0.21) (Randall and Selitto, 1957). There was also no significant difference between these two groups on the 50% mechanical withdrawal thresholds for fine filament von Frey hairs applied to the abdomen (Fig. 3F, *t*-test, *P* = 0.99).

**Figure 3.**
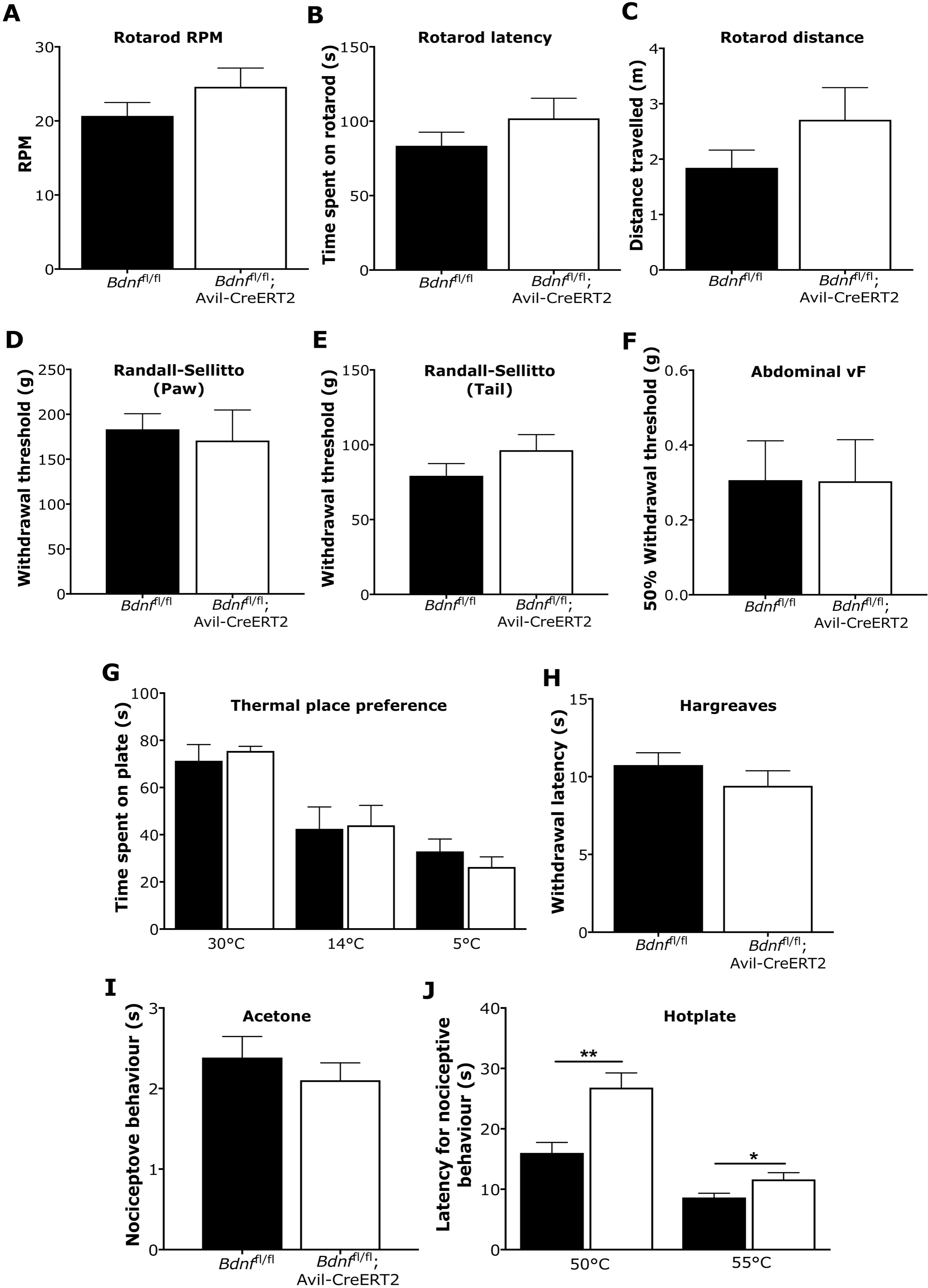
Motor function and acute pain behavior tests. (**A**) Motor function was accessed with Rotarod on speed (left panel), time spent on rotarod (middle panel) and distance walked on rotarod (right panel). (**B**) Mechanical pain threshold was examined with Randall-Sellitto apparatus on paw (left panel) and tail (right panel). (**C)** Mechanical light touch threshold were measured with von Frey hair on abdomen. (**D**) Cold pain was tested with thermal place preference at 30, 14 and 4 degrees. (**E**) Thermal pain was accessed with Hargreaves’ test. (**F**) Cold pain threshold was measured with acetone test. (**G**) Thermal pain behavior was examined with hot-plate test. Data were analysed with Student’s *t*-test (\**P* < 0.05, \*\**P* < 0.01).

Thermal place preference for innocuous warm (30°C), cool (14°C) and noxious cold (5°C) revealed no changes in thermal sensory function in the BDNF knockouts (Fig. 3G, *t-* test, *P* = 0.61, *P* = 0.91 and *P* = 0.36, respectively), and thermal pain thresholds in the Hargreaves’ test was comparable to littermate controls (Fig. 3H, *t-* test, *P* = 0.30) (Hargreaves *et al*, 1988). Acetone applied to the hind paw did not evoked significantly different pain behaviours from wildtype mice, also indicating no altered sensitivity to cold stimulation (Fig. 3I, *t*-test, *P* = 0.47). However, we did observe a hyposensitivity to noxious heat at temperatures of 50°C and 55°C in the hot plate test, which assess supraspinally mediated nociceptive behaviours (Fig. 3J, *t*-test, *P* = 002 and *P* = 0.03, respectively) (Woolfe and Macdonald, 1944). This mild thermal phenotype is similar to that we have previously reported with BDNF deletion from Na_v_1.8-expressing DRG neurons (Zhao *et al.*, 2006).

### Chronic pain models of inflammation, neuropathy and hyperalgesic priming

We used the intraplantar formalin test to assess effects of BDNF deletion from sensory neurons on inflammatory pain behaviour (Hunskaar *et al.*, 1985) (Fig. 4). *Bdnf*^fl/fl^; Avil-CreERT2 mice show comparable nocifensive behaviour to wildtype *Bdnf*^fl/fl^ mice in the first phase but show deficits in the second phase *(P* < 0.001, *t*-test). To determine the effects on evoked pain behaviour following neuropathy, we used the modified Chung model and partial sciatic nerve ligation model (Seltzer model). Following the modified Chung model of neuropathy (Fig. 5A), we observed prolonged mechanical hypersensitivity for the duration of 28 days in *Bdnf*^fl/fl^ control mice *(P* < 0.05 on Day 14, 21, 28 and *P* < 0.01 on Day 3, 5, 7 and 10, 1 way repeated measures ANOVA with Dunnett’s post-tests). In *Bdnf*^fl/fl^; Avil-CreERT2 mice, we observed an acute drop in mechanical withdrawal thresholds lasting up to 14 days *(P* < 0.05 on Day 5 and 14, *P* < 0.01 on Day 3, 7 and 10, 1 way repeated measures ANOVA with Dunnett’s post-tests), but began to see recovery from Day 5 followed by full recovery of withdrawal thresholds back to baseline in the chronic, later phases of the model at Day 21 and 28 (*P* > 0.05 on Day 21 and 28, 1 way repeated measures ANOVA with Dunnett’s post-tests). Following partial sciatic nerve ligation (Fig. 5B), we observed persistent mechanical hypersensitivity in both wildtype *Bdnf*^fl/fl^ mice and *Bdnf*^fl/fl^; Avil-CreERT2 mice *(P* < 0.05 and *P* < 0.01 in both groups at all measured time points, 1 way repeated measures ANOVA with Dunnett’s post-tests). Our combined data from the inflammatory formalin test and modified Chung model suggest that BDNF expressed in sensory neurons drives chronic, but not acute, nociceptive processes in inflammatory and some transection-related neuropathic pain.

**Figure 4.**
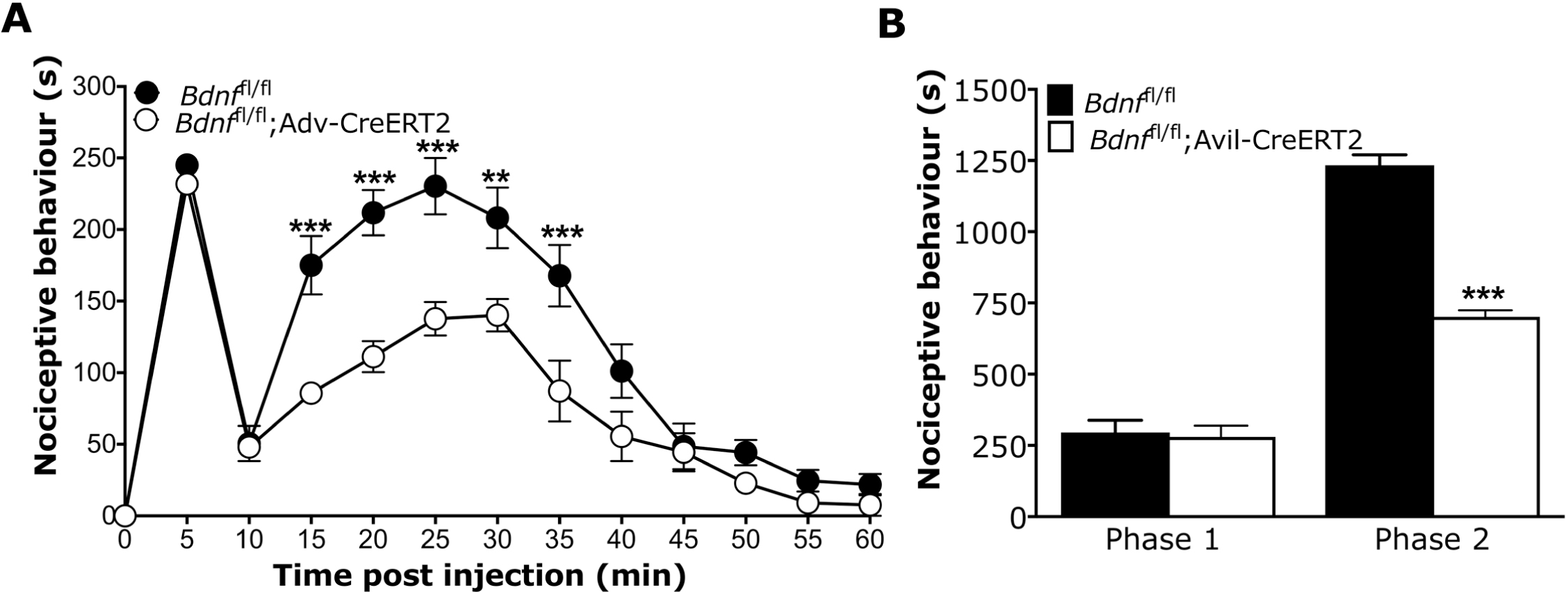
Formalin test. (**A**) Time course of formalin-induced nociceptive responses. (**B**) Phase I and Phase II summed nocicefensive behavior. A significant attenuation of pain behaviour in Phase 2 was observed in BDNF knockout mice. *Bdnf*^fl/fl^ (*n* = 6) and *Bdnf*^fl/fl^; Avil-CreERT2 (*n* = 6) were used. Data were analysed with either 2 way repeated measures ANOVA with Bonferroni post-tests (panel A, \*\**P* < 0.01, \*\*\**P* < 0.001) or Student’s *t-*test (panel B, \*\*\**P* < 0.001).

**Figure 5.**
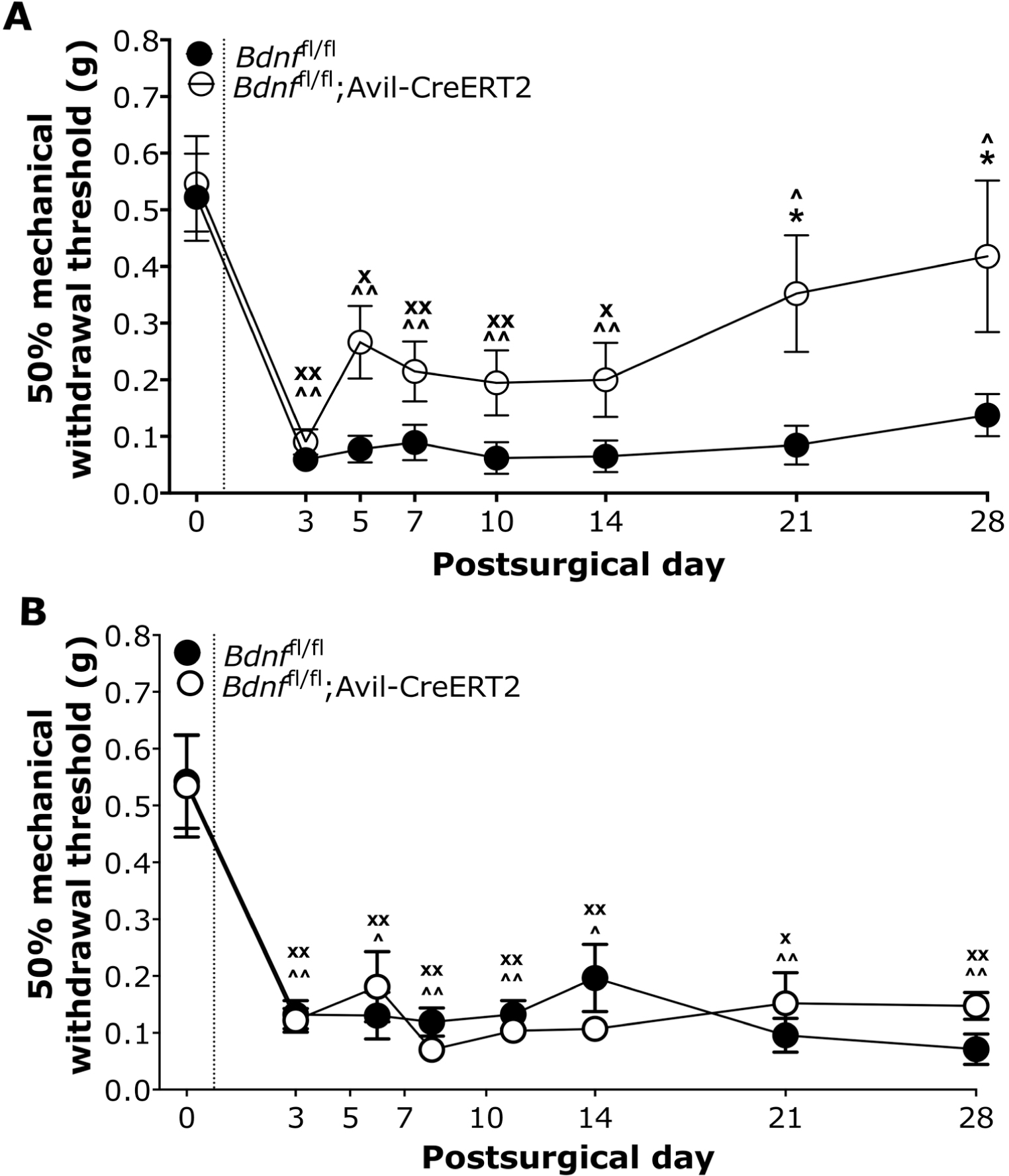
Neuropathic pain models. (**A**) A modified Chung surgical model was employed to assess development of neuropathic pain. (**B**) The Seltzer surgical model of neuropathy. *Bdnf*^fl/fl^ (*n* = 7) and *Bdnf*^fl/fl^; Avil-CreERT2 mice (*n* = 7) were tested in these two models. Data were analysed using 2 way repeated measures ANOVA with Bonferroni post-tests (\**P* < 0.05), 1 way repeated measures ANOVA with Dunnett’s post-tests for *Bdnf*^fl/fl^ group (^*P* < 0.05, ^*P* < 0.01), and 1 way repeated measures ANOVA with Dunnett’s post-tests for *Bdnf*^fl/fl^; Avil-CreERT2 group (x*P* < 0.05, x*P* < 0.01).

To further explore the distinct contribution of primary afferent-derived BDNF in acute and chronic nociceptive processing, we established a model of hyperalgesic priming to model the transition from acute to chronic pain in rodents, as described in other studies (Aley *et al.*, 2000; King *et al.*, 2012) (Fig. 6). Here, a prior injury can prime pain pathways to produce chronic pain following a subsequent insult. We first validated this model of hyperalgesic priming in C57BL/6J wildtype mice using an intraplantar priming injection of 25ul 1% carrageenan. This induces a transient mechanical hypersensitivity with a recovery to baseline thresholds within 72 hours (Fig. 6A). At Day 6, intraplantar injection of PGE2 leads to a short-lasting mechanical hypersensitivity in unprimed control mice, with recovery to baseline thresholds within one hour (Fig. 6A: 1 way repeated measures ANOVA with Dunnett’s post-tests, *P* < 0.001 at 30 minutes post-PGE2 in unprimed group). In contrast, primed mice develop long-lasting mechanical hypersensitivity lasting up to 7 days (Fig. 6A: 1 way repeated measures ANOVA with Dunnett’s post-tests, *P* < 0.001 at all time points post-PGE2 in primed group). We used this model to determine the contribution of BDNF to the transition from acute to chronic pain (Fig. 6B). We found that priming with intraplantar injection of carrageenan induced a transient mechanical hypersensitivity in both *Bdnf* conditional knockouts and littermate controls. Intraplantar injection of carrageenan alone is an established model of acute inflammation, and like our findings in the first phase formalin behaviour (Fig. 4), we observed no difference in nociceptive behaviour between BDNF knockouts and littermate controls. Littermate control mice recovered to baseline thresholds within 72 hours (Fig. 6B, 1 way repeated measures ANOVA with Dunnett’s post-tests, *P* < 0.001 up to 48 hours post-carrageenan in *Bdnf*^fl/fl^ group). However, we observed a significantly faster recovery to baseline thresholds in *Bdnf*^fl/fl^; Avil-CreERT2 mice (Fig. 6B, 1 way repeated measures ANOVA with Dunnett’s post-tests, *P<*0.001 up to 1 hour post-carrageenan in *Bdnf*^fl/fl^; Avil-CreERT2 group). At Day 6, mechanical withdrawal thresholds had returned to baseline values and effects of intraplantar injection of PGE2 were assessed. As demonstrated in our validation study in wildtype mice (Fig. 6A), primed *Bdnf*^fl/fl^ littermate controls developed prolonged mechanical hypersensitivity lasting up to 7 days post PGE2 (Fig. 6B, 1 way repeated measures ANOVA with Dunnett’s post-tests, *P<*0.001 up to day 7 post-PGE2 in *Bdnf*^fl/fl^ group). In contrast, primed *Bdnf*^fl/fl^; Avil-CreERT2 mice developed a shorter lasting hypersensitivity lasting only up to 60 minutes post-PGE2 (Fig. 6B, 1 way repeated measures ANOVA with Dunnett’s post-test, *P* < 0.001 up to 1 hour post-PGE2 in *Bdnf*^fl/fl^; Avil-CreERT2 group).

**Figure 6.**
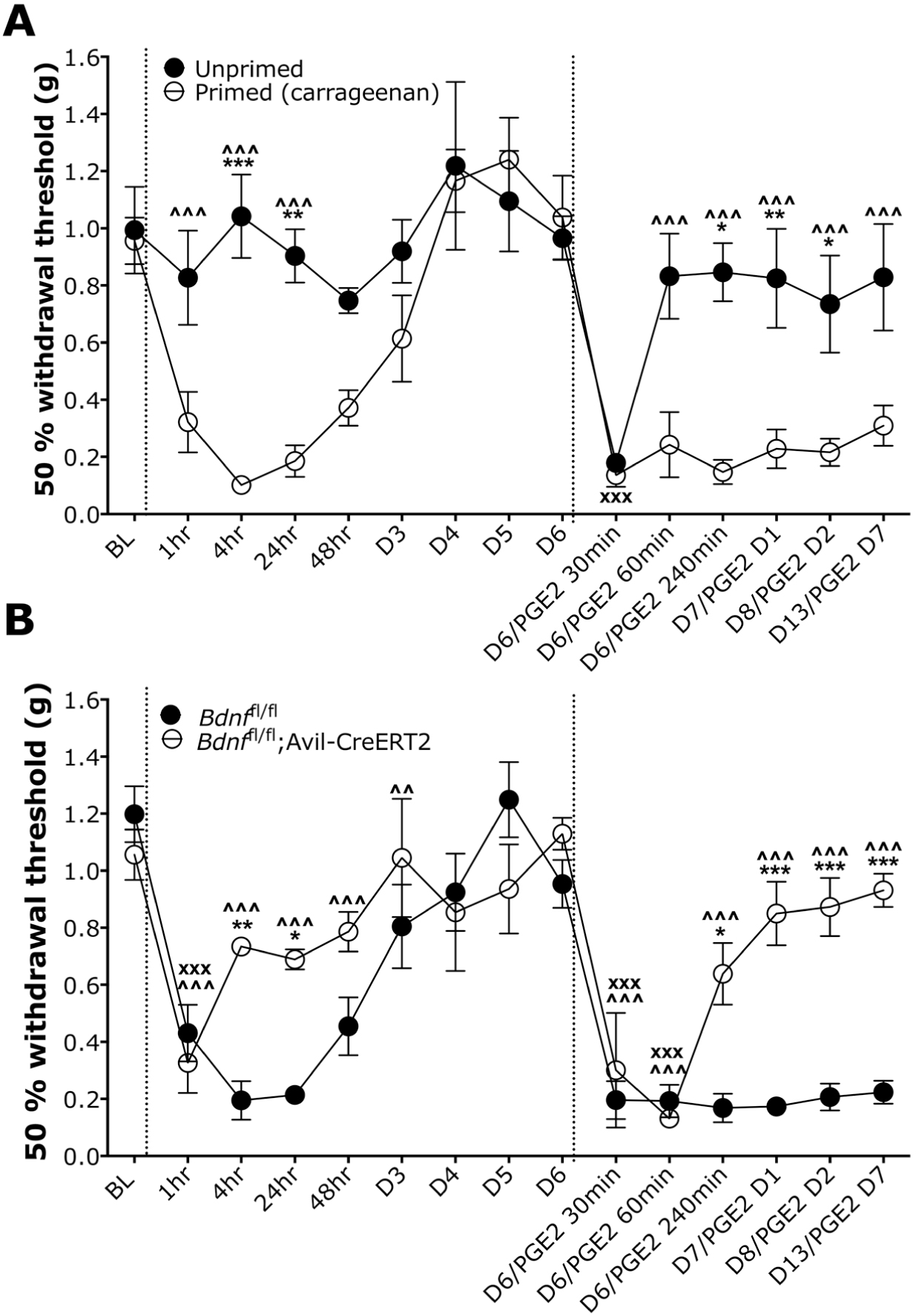
Hyperalgesic priming model. (**A**) a model of hyperalgesic priming with wild type C57BL/6J mice shows that priming mice with intraplantar carrageenan (first dotted line) confers prolonged mechanical hypersensitivity to PGE2 (second dotted line) compared to control unprimed mice (*n* = 6 in both groups). (**B**) *Bdnf*^fl/fl^; Avil-CreERT2 mice (*n* = 6) and their littermate controls *Bdnf^fl^*^/fl^ control mice (*n* = 7) develop transient mechanical hypersensitivity following the priming injection of intraplantar careegeenan, but BDNF mutant mice do not develop prolonged mechanical hypersensitivity following intraplantar PGE2, unlike their littermate controls. Data were analysed with 2 way repeated measures ANOVA with Bonferroni post-tests (\**P* < 0.05, \*\**P* < 0.01, \*\*\**P* < 0.001), 1 way repeated measures ANOVA with Dunnett’s post-tests for primed/BDNF^fl/fl^ groups (^^^*P* < 0.001), and 1 way repeated measures ANOVA with Dunnett’s post-tests for unprimed/*Bdnf*^fl/fl^; Avil-CreERT2 groups (xxx*P* < 0.001).

## Discussion

We deleted the *Bdnf* gene from adult peripheral sensory neurons to determine its contribution to acute and chronic pain processing. Our combined behavioural, electrophysiological and immunohistochemistry data show that BDNF released from sensory neurons does not significantly contribute to acute pain, but is necessary for the transition from acute to chronic inflammatory pain and some neuropathic pain states.

Tissue-specific gene ablation can provide important information about the relevance of a new drug targets, and time-specific gene deletion overcomes potential physiological compensatory mechanisms that can mask the true phenotypic contribution of a gene (Feil *et al.*, 1997; Metzger and Chambon, 2001). Advillin is a pan-neuronal marker of spinal and cranial sensory ganglia (Zurborg *et al.*; Marks *et al.*, 1998; Ravenall *et al.*, 2002), and we have previously generated BAC transgenic mice using the *Advillin* promoter to drive a tamoxifen-inducible CreERT2 recombinase construct that permits gene deletion in adult animals (Lau *et al.*, 2011). Here we used floxed *Bdnf* mice for Cre-mediated excision of *Bdnf* from sensory ganglia in order to determine the contribution of primary afferent-derived BDNF on the transition from acute to chronic pain. To assess the effects of *Bdnf* deletion from sensory neurons at the cell population level in DRG, we used immuno-staining with the neuronal markers neurofilament NF200 and peripherin. We found no difference in the total number of lumbar DRG neurons or proportions of large or small/medium diameter afferents following *Bdnf* deletion. This is in line with our previous findings that *Bdnf* deletion from Nav1.8-expressing neurons, an estimated 75% of all DRG neurons in lumbar ganglia (Shields *et al.*, 2012), also results a normal complement of DRG neurons (Zhao *et al.*, 2006).

To determine potential changes in spinal excitability we performed *in vivo* electrophysiological recordings from wide dynamic range neurons in the deep dorsal horn. Wide-dynamic range neurons in the deep dorsal horn receive converging primary afferent input, and their neural coding to peripheral stimulation conveys whether peripheral and central nociceptive processing has been significantly altered (Abrahamsen *et al.*, 2008; Bannister *et al.*, 2011; Sikandar *et al.*, 2013; O’Neill *et al.*, 2015). We measured evoked activity of spinal neurons to mechanical and thermal stimuli of low- and high-threshold intensities and found no significant impairment of graded coding to sensory stimulation in BDNF knockout mice (Fig. 2). We also found no difference in the overall amount of neuronal firing evoked by peripheral stimulation. These findings are in line with the normal neuronal complement of DRG neurons, indicating that peripheral input to the spinal cord is normal following *Bdnf* deletion from sensory neurons. Moreover, behavioural assays measuring acute reflexes to noxious stimuli showed no significant difference between BDNF knockout mice and littermate controls to mechanical, cold and most thermal assays (Fig. 3). However, the knockout mice were hyposensitive to the hotplate assay – this recruits bulbospinal reflexes, indicating that BDNF expression in sensory ganglia contributes to reflexes mediated from the brainstem. We have previously observed a similar hotplate phenotype following BDNF deletion from Na_v_1.8-expressing neurons (Zhao *et al.*, 2006).

Several lines of evidence support the role of enhanced DRG and spinal BDNF expression in persistent pain that follows injury-induced sensitisation of nociceptors (Cho *et al.*, 1997; Cho *et al.*, 1998; Lever *et al.*, 2003; Coull *et al.*, 2005; Li *et al.*, 2006; Lin *et al.*, 2011; Melemedjian *et al.*, 2013). However, the relative importance of BDNF released either from microglia or sensory neurons in chronic pain states is still unclear (Coull *et al.*, 2005; Zhuo *et al.*, 2011), and the contribution of microglial activity to persistent pain is further confounded by well established sex differences in immune-related nociceptive processing (Sorge *et al.*, 2011; Sorge *et al.*, 2015). In this study, we examine the role of primary afferent-derived BDNF on pain processing using both male and female mice.

To study the contribution of sensory BDNF to the development and maintenance of chronic pain, we used the formalin model of inflammation, a modified Chung and Seltzer models of neuropathy and a model of hyperalgesic priming. The formalin model entails biphasic pain behaviour – the first phase reflects acute, peripheral hypersensitivity and the second phase relates to chronic maintenance of pain through central sensitisation (Tjolsen *et al.*, 1992; Berge, 2011). We found comparable first phase nocifensive behaviour between BDNF knockouts and littermate controls, but significantly reduced second phase nocifensive behaviour in knockouts (Fig. 4). Similarly, we found that nerve-injury induced mechanical hypersensitivity was comparable to littermates in the acute phase following a modified Chung surgery, but BDNF knockout mice showed recovery of pain behaviour in a later chronic phase from 3 weeks after surgery (Fig. 5). Our data in the formalin and modified Chung surgery models suggest that BDNF signaling in sensory neurons is important for chronic, but not acute, pain processing.

Notably, we did not observe any difference in the development of mechanical hypersensitivity in the Seltzer model between BDNF knockout mice and littermate controls at any time point (Fig. 5B). Our observations support previous findings that the development of chronic pain in the modified Chung model is dependent on the expression of BDNF in sensory neurons, unlike partial sciatic nerve ligation (Obata *et al.*, 2006b). Previous studies have also reported differences in behavioural phenotypes (Kim *et al.*, 1997; Dowdall *et al.*, 2005) and in the contribution of distinct neuronal subsets across different rodent models of neuropathy (Minett *et al.*, 2012). Moreover, differences in levels of BDNF expression in DRG following distinct rhizotomy and transection models of neuropathy have been reported previously (Obata and Noguchi, 2006b; Obata *et al.*, 2006b).

We also validated a model of hyperalgesic priming to determine the role of sensory BNDF in the transition from acute to chronic pain. Hyperalgesic priming of the nociceptive system reflects long-lasting, latent hyper-responsiveness of nociceptors to inflammatory mediators subsequent to an inflammatory or neuropathic insult (Reichling and Levine, 2009). In rodents, repeated injections of algogenic substances with short-lasting acute effects can produce long-lasting hypersensitivity (Aley *et al.*, 2000; Parada *et al.*, 2003; Sluka *et al.*, 2003; Melemedjian *et al.*, 2014). Here we used intraplantar carrageenan as a priming agent (Fig. 6) to produce prolonged mechanical hypersensitivity to intraplantar PGE2 compared to animals that had not been primed. In *Bdnf*^fl/fl^ mice, we confirmed that priming with carrageenan precipitates long-lasting changes in PGE2-induction of pain behaviour. In contrast, we found that administration of intraplantar carrageenan in BDNF knockout mice produces an acute mechanical hypersensitivity, but does not prime mice to develop prolonged hypersensitivity to PGE2. Our data suggest that BDNF expression in sensory neurons mediates the transition from acute to chronic pain in a model of hyperalgesic priming. Other studies support our findings of this pro nociceptive role of BDNF in pain chronification, where spinal BDNF mediates prolonged PGE2 sensitivity in rodents primed with IL-6 (Melemedjian *et al.*, 2013) and sequestering BDNF in the cisterna magna can prevent IL-6-mediated hyperalgesic priming in a model of migraine (Burgos-Vega *et al.*, 2016).

An important implication of our findings is that BDNF expressed in sensory neurons is not essential for acute pain, but is critical for the transition from acute to chronic pain in models of inflammation, neuropathy and hyperalgesic priming. Studies of long term potentiation suggest that both pre- and post-synaptic release of BDNF regulates consolidation of late-LTP (Dean *et al.*, 2009; Jourdi *et al.*, 2009), and regulation of aPKCs at spinal synapses is likely to be a mechanism for BDNF to initiate and maintain chronic pain states (Melemedjian *et al.*, 2013). Different sources of BDNF may also drive nociceptive mechanisms in different preclinical pain models, i.e. the gender-specific contribution of microglial BDNF to chronic pain in a model of sciatic nerve cuffing (Coull *et al.*, 2005). A mechanistic discrepancy of neurotrophin signaling between rodent models likely underlies our findings that BDNF released from sensory neurons is essential for the development of chronic pain in a model of full, but not partial, nerve ligation (Obata *et al.*, 2006b).

In conclusion, our findings demonstrate the pro-nociceptive role of primary-afferent derived BDNF in mediating the transition from acute to chronic pain. These findings support the therapeutic potential of modulating BDNF for chronic pain syndromes (Nijs *et al.*, 2015b). Because BDNF expression is ubiquitously expressed in the nervous system, the development of targeted gene therapies for subsets of sensory neurons holds promise for providing adequate pain relief and overcoming side effects arising from central modulation (Glorioso and Fink, 2009; Fink *et al.*, 2011).

## Acknowledgements

We thank the Wellcome Trust, the MRC and the ARUK for funding this work. We also thank Dr James J. Cox for his comments on our manuscript.

## Funding

This work was supported by the Wellcome Trust (200183/Z/15/Z to J.Z. and J.N.W., and 101054/Z/13/Z to J.N.W.), the RCUK Medical Research Council (G091905 to J.N.W.) and the Arthritis Research UK (507928 GNRK G95 to J.N.W.)

